# Identification of anti-pathogenic activity among *in silico* predicted small molecule inhibitors of *Pseudomonas aeruginosa* LasR or Nitric Oxide Reductase (NOR)

**DOI:** 10.1101/2023.07.17.549273

**Authors:** Gemini Gajera, Niel Henriksen, Bryan Cox, Vijay Kothari

## Abstract

Antibiotic resistant *Pseudomonas aeruginosa* strains cause considerable morbidity and mortality. Identification of novel targets in this notorious pathogen is urgently warranted to facilitate discovery of new anti-pathogenic agents acting against it. Attacking non-essential targets is believed to be a potential anti-virulence strategy. This study attempted to identify small molecule inhibitors of two important proteins LasR and nitric oxide reductase (NOR) in *P. aeruginosa*. This bacterial pathogen possesses multiple quorum sensing (QS) systems to regulate expression of many of its genes including those associated with virulence. Among these QS systems, ‘Las’ system can be said to be the ‘master’ regulator, whose receptor protein is LasR. Similarly, NOR plays crucial role in detoxification of reactive nitrogen species. This study attempted *in silico* identification of potential LasR or NOR inhibitors through a virtual screen employing AtomNet^®^, a proprietary deep learning neural network. Following virtual screening of a large number of compounds for their affinity to LasR or NOR, a final subset of <100 compounds was created by clustering and filtering the top scoring compounds. These compounds were evaluated for their *in vivo* anti-pathogenic activity by challenging the model host *Caenorhabditis elegans* with *P. aeruginosa* in presence or absence of test compounds. Survival of the worm population in 24-well assay plates was monitored over a period of 5 days microscopically. Of the 96 predicted LasR inhibitors, 11 exhibited anti-*Pseudomonas* activity (23-96% inhibition of bacterial virulence as per third-day end point) at 25-50 µg/ml. Of the 85 predicted NOR inhibitors, 8 exhibited anti-*Pseudomonas* activity (40-85% inhibition of bacterial virulence as per second-day end point) at 25-50 µg/ml. Further investigation on molecular mode of action of active compounds is warranted.

## Introduction

Antimicrobial resistance (AMR) among infectious bacteria has emerged as a healthcare challenge of global concern [https://amr-review.org/]. The World Health Organization (WHO) has also published a global action plan on AMR in 2015. As per the Indian National Action Plan on Antimicrobial Resistance (NAP-AMR:2017 – 2021), India is among the nations with the highest burden of bacterial infections. The crude mortality from infectious diseases in India today is 417 per 100,000 persons. Situation in other developing countries is equally grave. Murray et al. (2022) estimated 1.27 million deaths attributable to bacterial AMR in 2019. Of the six leading pathogens identified by them responsible for maximum death toll, one is *Pseudomonas aeruginosa*. It is amongst the most notorious pathogenic bacteria, and its carbapenem-resistant phenotype has been listed by WHO among priority pathogens for development of new antibiotics [https://www.who.int/medicines/publications/WHO-PPL-Short_Summary_25Feb-ET_NM_WHO.pdf]. Antibiotic-resistant *P. aeruginosa* has been listed as an important pathogen by CDC (Centers for Disease Control, USA) (https://www.cdc.gov/drugresistance/biggest-threats.html) as well as DBT (Department of Biotechnology, India) (https://dbtindia.gov.in/sites/default/files/IPPL_final.pdf) against which new antimicrobials are urgently required.

The pipeline for new antibiotics does not contain sufficient number of promising candidates (Tacconelli et al., 2018). There is a dearth of novel antibacterial leads as well as targets (Singh and Barrett, 2006; Payne et al., 2007). Identification and validation of new targets in important pathogens is of utmost significance (Jaeger and Flohé, 2006; Ruparel et al., 2023). Conventional bactericidal antibiotics attack a very narrow range of targets in susceptible bacteria e.g., cell wall synthesis, protein synthesis, nucleic acid synthesis, etc. Since last decade there has been much interest in the research community regarding discovery of anti-virulence molecules which may attenuate the bacterial virulence without necessarily killing them. Such ‘pathoblockers’ are expected to compromise the ability of the pathogen to damage the host, by exerting their effect on non-essential targets such as bacterial quorum sensing (QS) (Groleau et al., 2020), stress-response machinery (Joshi and Kothari, 2022), metal homeostasis (Porcheron and Dozois, 2015; Hofmann et al., 2021) etc. The next-generation antibacterials preferably should be attacking such hitherto unexplored/ underexplored targets in pathogenic bacteria (Calvert et al., 2018; Kamal et al., 2018; Bonvicini et al., 2023).

Proper functioning of the chemical-signal based intercellular communication, known as quorum sensing (QS), is crucial for sufficient expression of virulence in pathogens like *P. aeruginosa*, as QS is an effective mechanism for regulating expression of multiple genes associated with a multitude of functions (Huang et al., 2020). Interrupting bacterial QS can be an effective strategy to combat pathogens. QS consists of two components, signal generation and signal response, respectively encoded by LuxI and LuxR homologues (Abisado et al., 2018; Della Sala et al., 2019). Inhibiting the “signal response” component of QS (e.g. LasR in *P. aeruginosa*) can notably compromise their ability to exert collective behaviour in response to environmental changes or host defence. Owing to the important role of Las system in overall QS circuit of *P. aeruginosa*, its receptor protein LasR, is believed to be a plausible anti-virulence target (Kostylev et al., 2019). LasR is a transcriptional activator of various virulence-associated genes in *P. aeruginosa*, which recognizes a specific signal molecule namely N-3-oxo-dodecanoyl-l-homoserine lactone (3O-C12-HSL) (Schuster et al., 2004). The LasR-3O-C12-HSL complex triggers the expression of multiple QS-regulated genes. Potential LasR inhibitors either may prevent its binding with its natural ligand or compromise its ability to affect expression of target genes (Maddocks, 2016). QS not being essential for bacterial survival, its inhibitors are expected to exert lesser selection pressure on bacterial population with respect to development of resistant phenotypes (Haque et al., 2018). Additionally, due to the overlap in QS systems among various gram-negative bacteria (Geske et al., 2008; Packiavathy et al., 2021), inhibitors effective against one gram-negative species may have broad-spectrum activity against multiple other gram-negative pathogens.

Pathogens striving to survive inside a host body are forced to face a variety of stresses such as iron deprivation, oxidative stress, nitrosative stress, etc. Bacteria employ antioxidant enzymes to counter reactive oxygen species, and similarly certain other enzymes to counter reactive nitrogen species (Poole, 2014). From the work done in our lab as well as literature survey, we consider the components of *P. aeruginosa* genome involved in responding to nitrosative stress to be potential targets. Among the components of nitrosative stress response in *P. aeruginosa*, one important enzyme is Nitric Oxide Reductase (NOR). This protein was one of the major targets of an anti-infective polyherbal formulation (*Panchvalkal*) investigated by us in recent past (Joshi et al., 2019; Ruparel et al., 2023). NOR also emerged among the top differentially expressed genes in *P. aeruginosa* treated by us with another anti-virulence polyherbal formulation-Enteropan (SRX15248092) or colloidal silver (SRX14392191) at sub-MIC levels.

NOR is an important detoxifying enzyme in *P. aeruginosa*, which is crucial to its ability to withstand nitrosative stress (e.g., in form of nitric oxide: NO). NOR has also been reported (Van Alst et al., 2007; Han et al., 2022) to be important for virulence expression of this pathogen, and thus can be a plausible target for novel anti-virulence agents. Molecules capable of inhibiting NOR can be expected to compromise the pathogen’s ability to detoxify NO, not allowing its virulence traits (e.g., biofilm formation, as NO has been indicated to act as a biofilm-dispersal signal) to be expressed fully (Toyofuku and Yoon, 2018). The test molecules capable of inhibiting NOR may emerge as novel anti-biofilm agents not only against *P. aeruginosa* but against multiple pathogens as NO is reported to be perceived as a dispersal signal by various gram-negative and gram-positive bacteria (Barraud et al., 2015). Thus, NOR inhibitors may be expected to have a broad-spectrum activity against multiple pathogens. Major function of NOR is to detoxify NO generated by nitrite reductase (NIR). Nitric oxide (NO) is a toxic byproduct of anaerobic respiration in *P. aeruginosa*. NO-derived nitrosative species can damage DNA, and compromise protein function. Intracellular accumulation of NO is likely to be lethal for the pathogen (Carvalho et al., 2022). It can be logically anticipated that *P. aeruginosa*’s ability to detoxify NO will be compromised under the influence of a potent NOR inhibitor. Since NO seems to have a broad-spectrum anti-biofilm effect, NOR activity is essential for effective biofilm formation by the pathogens. NOR activity and NO concentration can modulate cellular levels of c-di-GMP, which is a secondary messenger molecule recognized as a key bacterial regulator of multiple processes such as virulence, differentiation, and biofilm formation (Wang et al., 2022). In the mammalian pathogens, the host’s macrophages are a likely source of NO. NOR expressed by the pathogen provides protection against the host defence mechanism. Since NOR activity is known to be important in multiple pathogenic bacteria (e.g., *P. aeruginosa, Staphylococcus aureus, Serratia marcescens*) for biofilm formation, virulence expression, combating nitrosative stress, and evading hose defense; NOR seems to be an important target for novel anti-pathogenic agents. Any molecule capable of interfering with bacterial NOR activity is likely to be an effective anti-pathogenic agent, since bacterial populations require NOR for various purposes including detoxification and evasion of host defences (Kakishima et al., 2007). A potential NOR inhibitor besides troubling the pathogen directly, may also boost its clearance by the host macrophages.

This study aimed at screening 96 compounds identified through a virtual screen as potential LasR inhibitors, and 85 compounds predicted to be NOR inhibitors *in silico* for their possible anti-virulence activity against *P. aeruginosa* in the model host *Caenorhabditis elegans*. Any such potent NOR or LasR inhibitors identified through this study may prove to be useful lead(s) for novel anti-pathogenic drug development. They can be useful either as standalone therapy or in combination with conventional antibiotics.

## Methods

### Virtual Screening

The virtual screening from a library of approximately 3 million compounds was conducted using AtomNet^®^ screening platform (Wallach et al., 2015). AtomNet^®^ is a proprietary deep learning neural network useful for structure-based drug design and discovery through its small molecule binding affinity prediction capacity.

### Screening for LasR binding ligands

There are a number of available crystal structures of LasR in complex with small molecules, including the endogenous ligand and other agonists. Although discovery of a potent antagonist is preferable, the virtual screen attempted to find novel chemical matter which binds at the desired site and the mode of binding may be analysed later. All of the available crystal structures were considered as receptor templates for virtual screening. The highest resolution structure, PDB 3IX3, chosen as the ligand binding pocket is deep and solvent excluded and appears well-suited for binding small molecules (Figure 1). The screening volume was restricted to the binding pocket surrounding the existing ligand. A screening library of approximately 3 million compounds was exhaustively pose-sampled and scored, followed by filtering for drug-like properties, and selection of the top 96 compounds for ordering in physical form.

**Figure 1.**
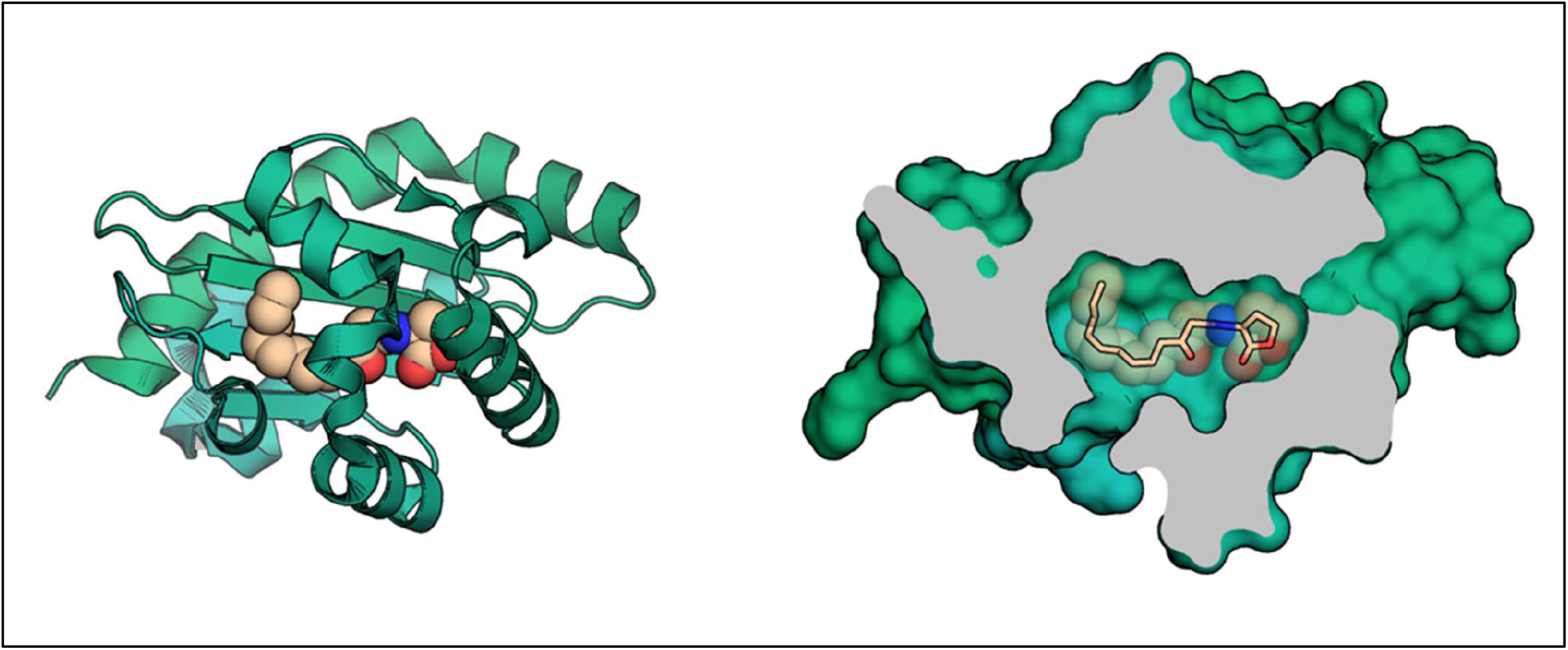
The structure of LasR (green) in complex with its natural ligand 3O-C12-HSL (tan). Coordinates taken from PDB 3IX3.

### Screening for NOR binding ligands

The structure of *Pseudomonas aeruginosa* NOR bound to an antibody fragment has been determined by crystallography (PDBID 3O0R). This structure reveals NOR composed of two subunits: NORB containing 12 transmembrane helices and NORC with a transmembrane helix and hydrophilic periplasmic domain. Three heme cofactors are complexed in NOR. Two hemes are deep in the NORB transmembrane region, and the third heme spans the NORB-NORC interface. No drug-like small molecule inhibitors have been reported for NOR, and no druggable pocket is obvious from the reported structure. To identify potential targetable sites, the structure of NOR was analyzed by fPocket. This analysis revealed a pocket at the interface of NORB and NORC and near one of the NORB heme cofactors. This positioning suggests that small molecules binding to this pocket may disrupt norB-norC interactions and/or inhibit heme-mediated electrochemistry. The virtual screen therefore seeked to identify small molecules that bind to this pocket in NOR and potentially disrupt enzymatic function. A library of approximately three million compounds was screened for identifying compounds capable of virtual binding at the selected target site on NOR (Figure 2) using AtomNet^®^. Top scoring compounds were clustered and filtered to arrive at a final subset of 85 deliverable compounds.

**Figure 2:**
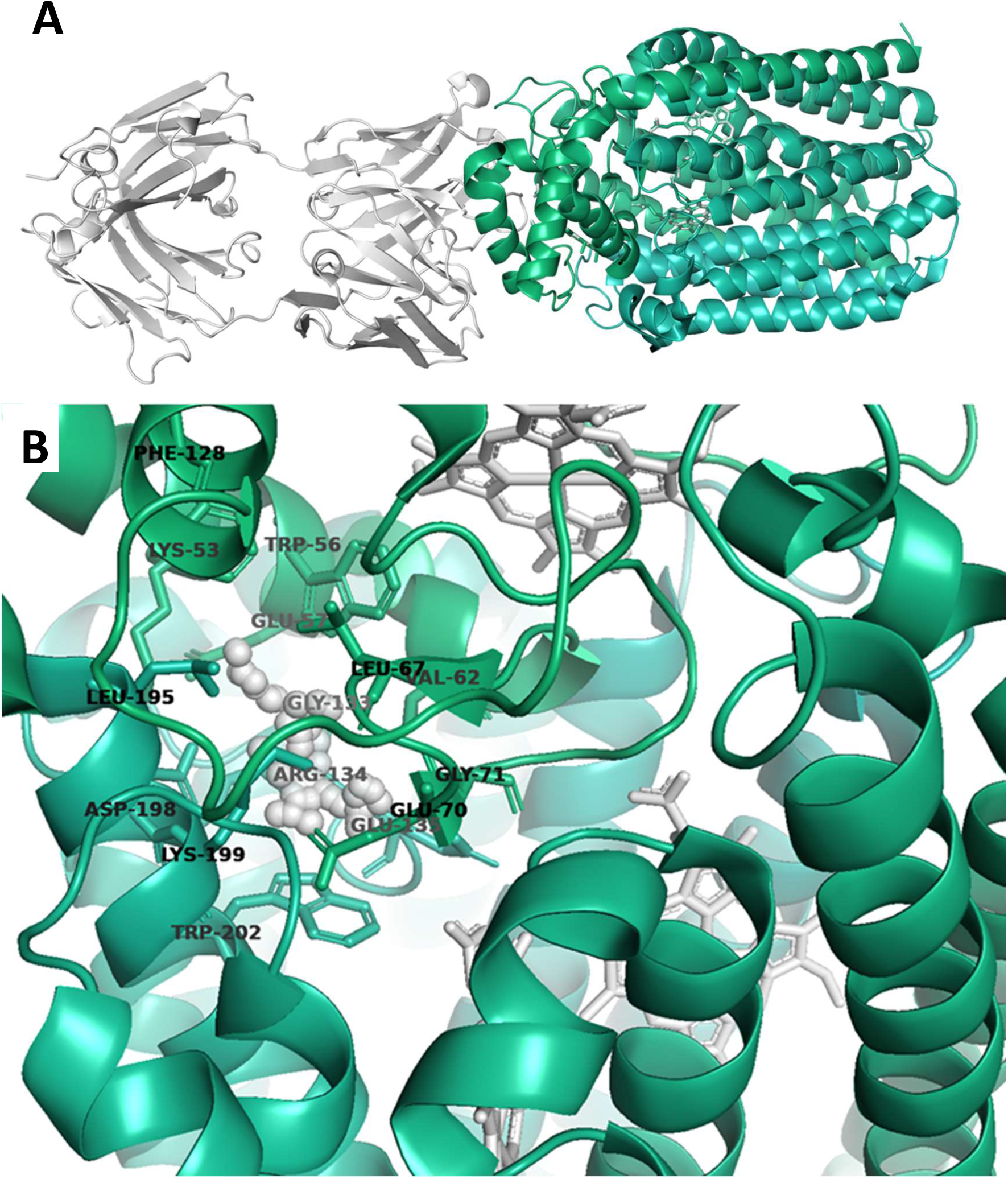
Nitric Oxide Reductase (NOR). (A) Crystal structure of NOR complexed with an antibody fragment (white cartoons) and with three heme cofactors (white sticks). (B) Proposed target for virtual screen (white spheres) with surrounding residues from NOR chains B and C showing the proximity to the heme cofactors.

### Test Compounds

Ninety six compounds showing *in silico* affinity to LasR (Table S1), and 85 compounds showing *in silico* affinity to NOR (Table S2) were ordered in physical form from Enamine (Kyiv, Ukraine) and Mcule (Budapest, Hungary) respectively. Test compounds were stored in refrigerator, and reconstituted in DMSO (500-1000 µl) (Merck) on the day of assay.

### Bacterial strain

The *P. aeruginosa* strain used in this study was sourced from our internal culture collection, which has been characterized by us for its antibiotic susceptibility/ resistance, pigment production and certain other virulence traits. Its antibiogram (https://www.biorxiv.org/content/10.1101/2023.06.27.546803v1.supplementary-material) generated through a disc-diffusion assay performed as per NCCLS guidelines revealed it to be resistant to 8 antibiotics (co-trimoxazole, augmentin, nitrofurantoin, ampicillin, chloramphenicol, clindamycin, cefixime, and vancomycin) belonging to 5 different classes. Hence it can be described as a multidrug resistant (MDR) strain. As reported in our earlier publications (Joshi et al., 2019; Patel et al., 2019) with this strain, it is a haemolytic strain capable of producing the QS-regulated pigments (pyocyanin and pyoverdine), and also of biofilm formation. We maintained this bacterium on Pseudomonas agar (HiMedia). While culturing the bacteria for *in vivo* assay, they were grown in Pseudomonas broth (magnesium chloride 1.4 g/L, potassium sulphate 10 g/L, peptic digest of animal tissue 20 g/L, pH 7.0 ± 0.2).

### Nematode host

*C. elegans* (N2 Bristol) was used as the model host in this study. Worms were maintained on NGM [Nematode Growing Medium; 3 g/L NaCl (HiMedia, MB023-500G), 2.5 g/L peptone (HiMedia), 17 g/L agar-agar (HiMedia), 1 M CaCl_2_ (HiMedia), 1 M MgSO_4_ (Merck), 5 mg/mL cholesterol (HiMedia), 1 M phosphate buffer of pH 6,] agar plate with *E. coli* OP50 (LabTIE B.V., the Netherlands) as food. For synchronization of the worm population, adult worms from a 4–5 days old NGM plate were first washed with distilled water, and then treated with one mL of bleaching solution [Water + 4% NaOCl (Merck) +1N NaOH (HiMedia) + in 3:1:1 proportion], followed by centrifugation (1500 rpm at 22°C) for 1 min. Eggs in the resultant pellet were washed multiple times with sterile distilled water, followed by transfer onto a new NGM plate seeded with *E. coli* OP50. L3-L4 stage worms appearing on this plate after 2–3 days of incubation at 22°C were used for further experimentation.

### Virulence assay

*P. aeruginosa* grown in Pseudomonas broth (at 35±1°C for 21 h with intermittent shaking) was allowed to attack *C. elegans* (L3-L4 stage) in a 24-well plate (HiMedia) in presence or absence of test compounds, and their capacity to kill the worm population was monitored over a period of 5 days. In each well, there were 10 worms in M9 buffer (3g/ L KH_2_PO_4_; 6g/L Na_2_HPO_4_; 5 g/L NaCl), which were challenged with *P. aeruginosa* by adding 100 µL (OD_764_=1.30) of bacterial culture grown in Pseudomonas broth. Appropriate controls i.e. worms exposed neither to test compound nor bacteria; worms exposed to test compound, but not to bacterial pathogens (toxicity control); worms challenged with bacteria in presence of 0.5%v/v DMSO (vehicle control); and worms challenged with bacteria in presence of 0.5 µg/mL ofloxacin (positive control) were also included in the experiment. Incubation was carried out at 22°C. Number of dead vs. live worms were counted every day for 5 days by putting the plate with lid under a light microscope 4X objective. Straight non-moving worms were considered to be dead. Plates were gently tapped to confirm lack of movement in the apparently-dead worms. On the last day of the experiment, when plates could be opened, their death was also confirmed by prodding them with a straight wire, wherein no movement was taken as confirmation of death.

## Statistics

Results reported are means of three replicates. Statistical significance was assessed through a t-test performed using Microsoft Excel. P-values ≤0.05 were considered to be significant.

## Results

### Anti-pathogenic activity of potential LasR inhibitors

Results of anti-virulence assay for all active compounds are presented in Figure-3. Since *P. aeruginosa* strain used by us could kill all worms within 18-36 h, any end-point beyond that can be taken as valid for labelling any compound as ‘active’ or ‘inactive’. However, to have more robust interpretation, we continued worm counting in assay plates till five days for comparing number of live worms in experimental vs. control wells.

**Figure 3.**
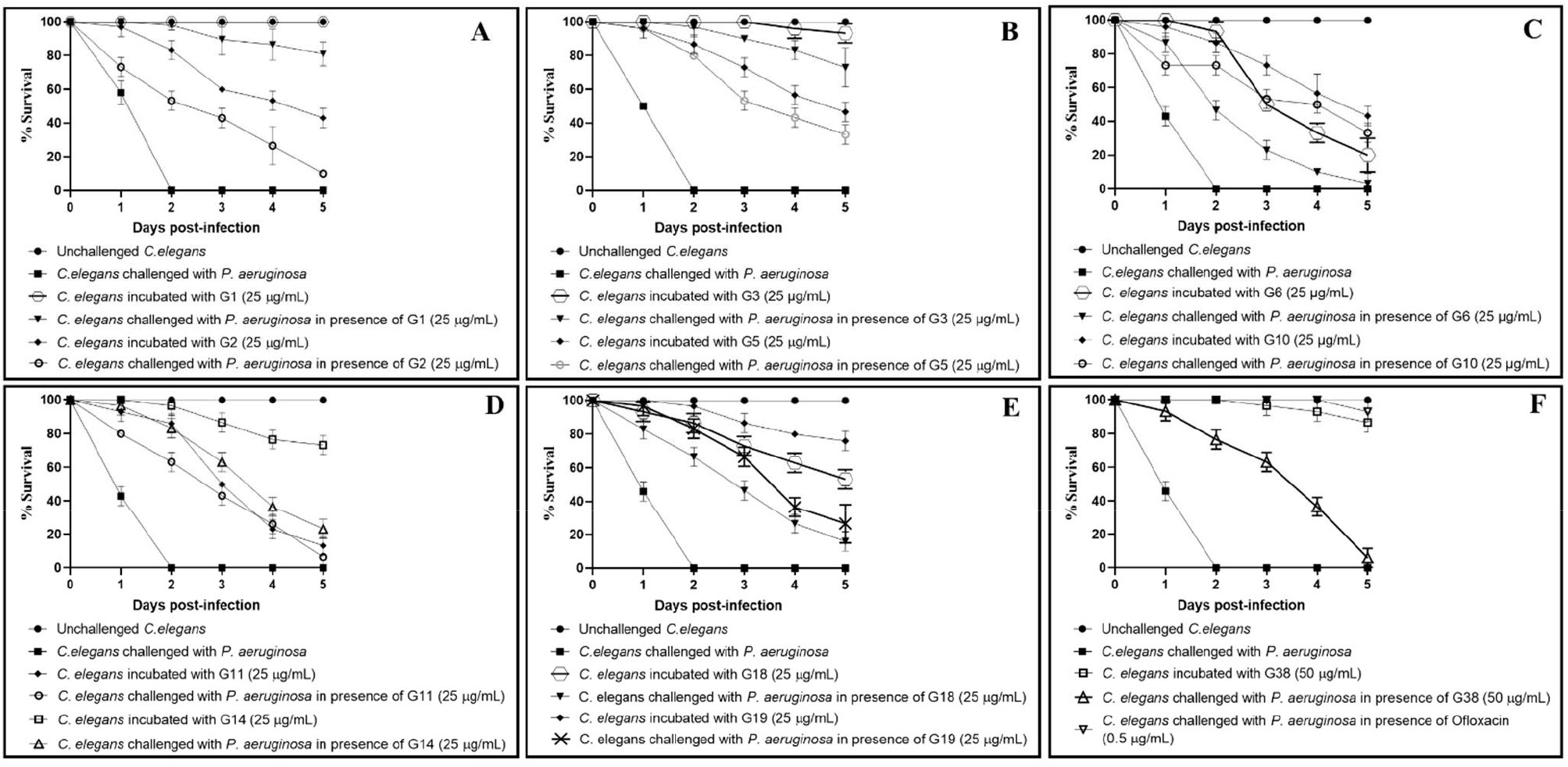
*P. aeruginosa*’s virulence towards the host worm gets attenuated in presence of certain predicted LasR inhibitors. **(A)** *P. aeruginosa* could kill 90%±9.19** and 43%±5.7*** lesser worms in presence of G1 (Z30981775) and G2 (Z65195564) respectively. **(B)** *P. aeruginosa* could kill 90%±0*** and 53%±5.7*** lesser worms in presence of G3 (Z212728858) and G5 (Z354444420) respectively. **(C)** *P. aeruginosa* could kill 23%±5.7*** and 53%±5.7*** lesser worms in presence of G6 (Z400859658) and G10 (Z1426094174) respectively. **(D)** *P. aeruginosa* could kill 43%±5.7*** and 63%±5.7*** lesser worms in presence of G11 (Z1625994950) and G14 (Z104586200) respectively. **(E)** *P. aeruginosa* could kill 46%±5.7*** and 67%±5.7*** lesser worms in presence of G18 (Z1084397894) and G19 (Z1212781307) respectively. **(F)** *P. aeruginosa* could kill 63.3%±5.7*** and 100%±0*** lesser worms in presence of G38 (Z89293640) and ofloxacin (0.5 µg/mL) respectively. Later was employed as a positive control at its sub-MIC level, and it did allow progeny formation in worm population third day onward. DMSO (0.5%v/v) present in the ‘vehicle control’ neither affected virulence of the bacterium towards *C. elegans*, nor did it show any effect on worm survival. **p<0.01, ***p<0.001. The percent values reported pertain to the worm survival 3-days post infection.

Eleven of the 88 DMSO-soluble compounds (i.e. 12.5%) assayed exhibited anti-pseudomonas activity (23-96% as per third-day end point) at 25-50 µg/mL (Table-1). These 11 compounds (G1, G2, G3, G5, G6, G10, G11, G14, G18, G19 and G38) should be tested at still lesser concentrations to find out minimum effective concentration (MEC). Eight of the test compounds were found to possess dual activity i.e., anti-pathogenic as well as anthelmintic. Eight of the active anti-Pseudomonas compounds (G2, G5, G6, G10, G11, G14, G18, G19) identified in this study were also toxic to the host worm at concentrations employed. Hence, they should be tested at still lower concentrations. It is possible that their lower concentrations may exhibit anti-pathogenic activity without exerting any toxicity towards the eukaryotic host. Masking of the anti-pathogenic activity by anti-worm activity of the same compound (Table-2) needs to be paid attention while interpreting the results.

**Table 1.** List of predicted LasR inhibitors found to possess *in vivo* anti-*P. aeruginosa* activity.

| Lab Code | Manufacturer's Code | Conc.<br>($\mu\text{g/mL}$) | % reduction in bacterial virulence | |
| --- | --- | --- | --- | --- |
|  |  |  | 3<sup>rd</sup> day end-point | 5<sup>th</sup> day end-point |
| G1 | Z30981775 | 25 | 90±9.19** | 82±7.07*** |
| G2 | Z65195564 | 25 | 43±5.7*** | 10±0*** |
| G3 | Z212728858 | 25 | 90±0*** | 73±11.5*** |
| G5 | Z354444420 | 25 | 53±5.7*** | 33±5.7*** |
| G6 | Z400859658 | 25 | 23±5.7*** | 3±5.7 |
| G10 | Z1426094174 | 25 | 53±5.7*** | 33±5.7*** |
| G11 | Z1625994950 | 25 | 43±5.7*** | 6±11.5 |
| G14 | Z104586200 | 25 | 63±5.7*** | 23±5.7*** |
| G18 | Z1084397894 | 25 | 46±5.7*** | 16±5.7** |
| G19 | Z1212781307 | 25 | 67±5.7*** | 26±11.5** |
| G38 | Z89293640 | 50 | 63.3±5.7*** | 6±5.7 |

\*\*p<0.01, \*\*\*p<0.001

**Table 2.** Anti-pathogenic activity of potential LasR inhibitors may be masked by their anthelmintic activity.

| Lab Code | Manufacturer's Code | Conc (µg/mL) | % anti-pathogenic activity based on fifth day end-point |  |
| --- | --- | --- | --- | --- |
|  |  |  | without nullifying compound's toxicity towards worms | After nullifying compound's toxicity towards worms |
| G2 | Z65195564 | 25 | 10±0*** | 67±0*** |
| G5 | Z354444420 | 25 | 33±5*** | 87±5*** |
| G6 | Z400859658 | 25 | 3±5.7 | 83±5.7*** |
| G10 | Z1426094174 | 25 | 33±5*** | 90±5*** |
| G11 | Z1625994950 | 25 | 6±11.5 | 93±11.5*** |
| G14 | Z104586200 | 25 | 23±5.7** | 50±5.7*** |
| G18 | Z1084397894 | 25 | 16±5.7** | 63±5.7*** |
| G19 | Z1212781307 | 25 | 26±11.5** | 53±11.5*** |
| G38 | Z89293640 | 50 | 6±5.7 | 20±5.7** |

\*\*p<0.01, \*\*\*p<0.001

Further, *in vitro* incubation of bacteria with these compounds is required to find out whether these compounds exhibit bactericidal / bacteriostatic/ anti-virulence activity. While evaluating any compound(s) for their anti-virulence activity, it should be kept in mind that even compounds capable of curbing bacterial virulence partially can be potentially useful in combination with conventional antibiotics. Such compounds may be potential resistance-modifiers. Even as a standalone therapy, they may be of indirect help to host immune system by reducing the overall bacterial load to be cleared by the immune system (Walsh, 2003; Allen et al., 2014). Three of these anti-pathogenic compounds (G1, G3, G38) did not exhibit any notable toxicity towards the host worm, and hence seem to be the most logical candidates for further investigation. These compounds should be tested against multiple species of antibiotic-resistant bacteria to know whether they are broad-spectrum antimicrobials. Additionally, their effect on bacterial gene expression at whole transcriptome level should also be investigated to elucidate the underlying molecular mechanisms.

### Anti-pathogenic activity of potential NOR inhibitors

Of the total 85 compounds received, 10 were insoluble in the vehicle solvent DMSO. Remaining 75 compounds were assayed for their possible anti-pathogenic activity by challenging the host worm with *P. aeruginosa* in presence or absence of test compounds. Eight (∼11% of the all compounds tested) of the test compounds were able to rescue the worm population from the pathogen-induced death by 40-85% (second day end-point) (Fig-4; Table-3).

**Figure 4.**
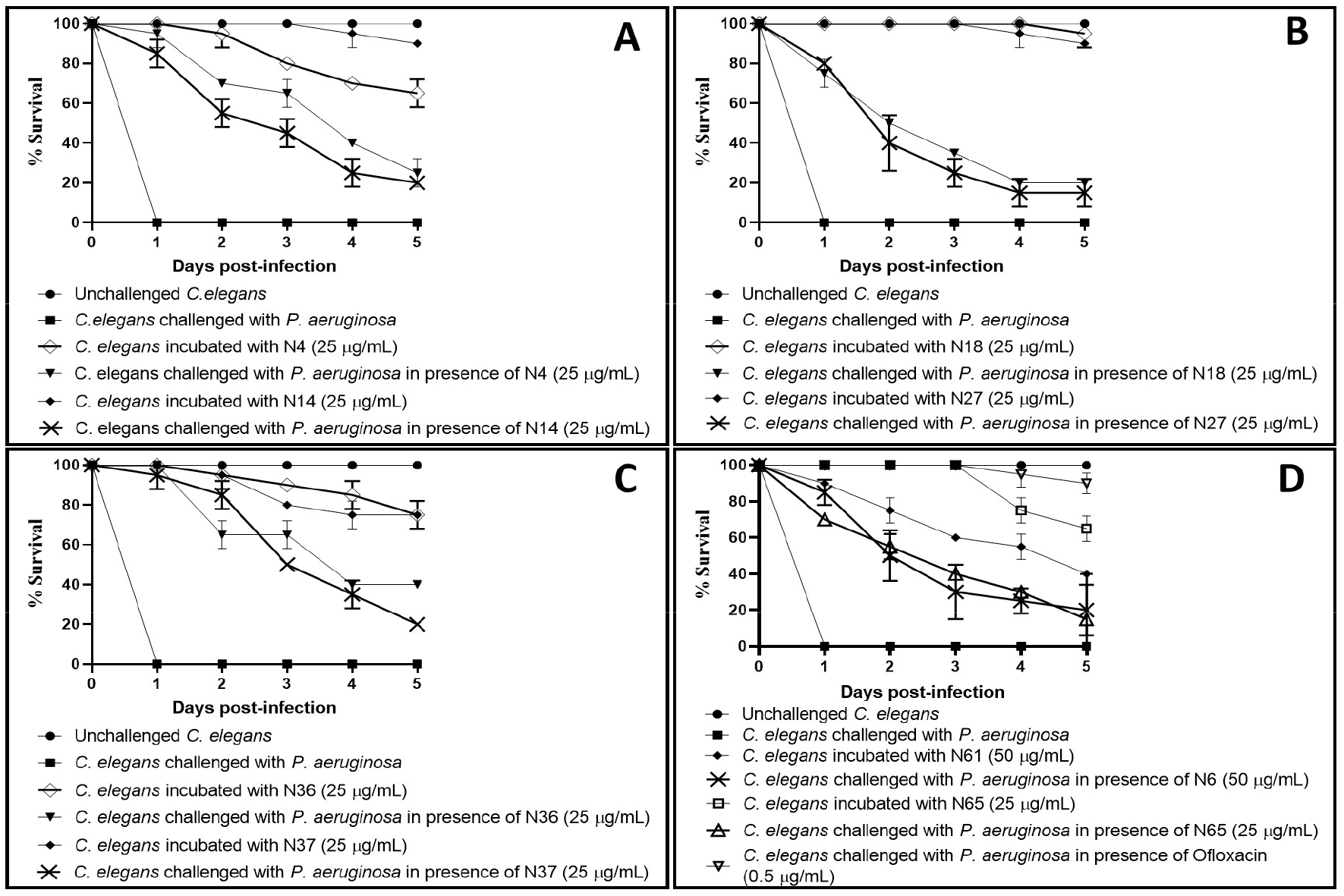
*P. aeruginosa*’s virulence towards the host worm gets attenuated in presence of certain predicted NOR inhibitors. **(A)** *P. aeruginosa* could kill 70%±0*** and 55%±7**lesser worms in presence of N4 (Z954454636) and N14 (Z1765101069) respectively. **(B)** *P. aeruginosa* could kill 50%±0*** and 40%±14* lesser worms in presence of N18 (Z110018576) and N27 (Z1611882500) respectively. **(C)** *P. aeruginosa* could kill 65%±7* and 85%±7** lesser worms in presence of N36 (Z397755956) and N37 (Z1190350270) respectively. **(D)** *P. aeruginosa* could kill 50%±14**, 55%±0.7** and 100%±0*** lesser worms in presence of N61 (Z2740017161), N65 (Z1874308288) and ofloxacin (0.5 µg/mL) respectively. Later was employed as a positive control at its sub-MIC level, and it did allow progeny formation in worm population third day onward. DMSO (0.5%v/v) present in the ‘vehicle control’ neither affected virulence of the bacterium towards *C. elegans*, nor did it show any effect on worm survival. **p<0.01, ***p<0.001. The percent values reported pertain to the worm survival 2-days post infection.

**Table 3.** List of predicted NOR inhibitors found to possess *in vivo* anti-*P. aeruginosa* activity.

| Lab Code | Manufacturer's Code | Conc.<br>(µg/mL) | % reduction in bacterial virulence |  |
| --- | --- | --- | --- | --- |
|  |  |  | 1<sup>st</sup> day end-point | 2<sup>nd</sup> day end-point |
| N4 | Z954454636 | 25 | 95±7*** | 70±0*** |
| N14 | Z1765101069 | 25 | 85±7** | 55±7** |
| N18 | Z110018576 | 25 | 75±7** | 50±0*** |
| N27 | Z1611882500 | 25 | 80±0*** | 40±14* |
| N36 | Z397755956 | 25 | 100±0*** | 65±7* |
| N37 | Z1190350270 | 25 | 95±7** | 85±7** |
| N61 | Z2740017161 | 50 | 85±7** | 50±14** |
| N65 | Z1874308288 | 25 | 70±0*** | 55±0.7** |

\*\*p<0.01, \*\*\*p<0.001

Five of the test compounds were found to possess dual activity i.e., anti-pathogenic as well as anthelmintic. Five of the active anti-Pseudomonas Compounds (N4, N36, N37, N61, N65) identified in this study were also toxic to the host worm at concentrations employed. Hence, they (excluding N61) should be tested at still lower concentrations. It is possible that their lower concentrations may exhibit anti-pathogenic activity without exerting any toxicity towards the eukaryotic host. Masking of the anti-pathogenic activity by anti-worm activity of the same compound (Table-4) needs to be paid attention while interpreting the results.

**Table 4.** Anti-pathogenic activity of potential NOR inhibitors may be masked by their anthelmintic activity.

| **Lab Code** | **Manufacturer's Code** | **Conc (µg/mL)** | **% anti-pathogenic activity based on fifth day end-point** |  |
| --- | --- | --- | --- | --- |
|  |  |  | **without nullifying compound's toxicity towards worms (A)** | **After nullifying compound's toxicity towards worms (B)** |
| N4 | Z954454636 | 25 | 25±7** | 60±7** |
| N36 | Z397755956 | 25 | 40±0** | 65±0*** |
| N37 | Z1190350270 | 25 | 20±0*** | 45±0*** |
| N61 | Z2740017161 | 50 | 20±14 | 80±14** |
| N65 | Z1874308288 | 25 | 15±21 | 40±21** |

\*\*p<0.01, \*\*\*p<0.001

### Conclusion

This study is a preliminary demonstration of the utility of virtual screening approach for discovery of potentially novel anti-pathogenic agents. Virtual screening can reduce the number of compounds required to be actually subjected to wet-lab assays, and thus reducing the investment of labour, time, and money. Among the top 181 compounds predicted through virtual screening to be capable of binding to NOR or LasR of *P. aeruginosa*, we could detect *in vivo* anti-*P. aeruginosa* activity in 19 (i.e. 10.4% of all compounds tested) of them in the model host *C. elegans*. As per our search on PubChem on June 1, 2023, these 19 compounds have yet not been reported to possess any kind of biological activity, and hence we believe this to be the first report of anti-pathogenic activity in these compounds. Further investigation on these active compounds with respect to their mode of action is warranted, which besides confirming their antibacterial activity will also provide additional validation to the targetability of NOR and LasR.

## Limitations

The anti-virulence assay performed in this study is not specific to LasR or NOR, hence precise mode of action of active anti-pathogenic compounds warrants further confirmatory assays. We could not carry out *in vitro* MIC/MBC assay for active compounds owing to limited quantity at our disposal, hence it was not possible to distinguish between growth-inhibitory and virulence-inhibitory (i.e. non-antibiotic action) activity. Further, anti-pathogenic activity of some of the compounds might be masked by their anthelmintic activity (Table 2-4).

## Acknowledgement

Authors thank Nirma Education and Research Foundation (NERF), Ahmedabad for infrastructural support.

